# A meta-analysis of age-dependent changes in extracellular vesicle proteins in *C. elegans*

**DOI:** 10.1101/2024.11.28.625880

**Authors:** Prasun Kumar Bhunia, Prasad Kasturi

## Abstract

Extracellular vesicles (EVs) contribute to the maintenance of organism-wide proteostasis by mediating intercellular communication. Loss of proteostasis and altered intercellular communication are associated with aging and age-related diseases, suggesting key roles for EVs. However, it is unclear how the proteome of the EVs changes with age. To identify EV-associated proteins (EVAPs) and their fate with age, we curated publicly available proteome data from *C. elegans* model organism. Our analysis reveals that EVs carry proteins that involve protein quality control. We found that abundance of the EV proteins changes significantly with age. Many of these EV proteins also aggregate with age and overlap with Aβ driven protein aggregates. We also observe that a subset of proteins that alter their abundance in response to heat stress and pathogen infections are also associated with the EVs. Further, we identify human orthologs of *C. elegans* EVAPs from human brain tissues affected with Alzheimer’s disease. This meta-analysis highlights EVs proteome composition, their abundance changes, and aggregation during aging, disease and stress conditions. Overall, this study provides new insights into the dynamics of EV proteins during aging and may possibly help in identifying potential biomarkers for age-related diseases.

## 1. Introduction

Aging is a complex, multifactorial process characterized by the gradual decline in the cellular functions, leading to increased susceptibility to age-related diseases and ultimately, death. One of the hallmarks of aging is loss of protein homeostasis or proteostasis (López-Otín et al., 2023), the state of balanced protein synthesis, folding, and degradation (Hipp et al., 2019). While the molecular chaperones assist in protein folding and prevent aggregation, the proteasomes and autophagy systems degrade misfolded or damaged proteins. As organisms age, the ability to maintain proteostasis deteriorates, leading to the accumulation of damaged or misfolded proteins, which is a hallmark of many age-associated diseases, including neurodegenerative disorders and cancer (Wilson et al., 2023). In an organism-wide context, proteostasis is not limited to individual cells or tissues, but requires coordination across the entire organism. This coordination is facilitated by inter-tissue communication, a process through which tissues and organs exchange signals to maintain systemic balance (Sala et al., 2017; Morimoto 2020). Notably, extracellular vesicles (EVs) have emerged as critical mediators of inter-tissue communication.

Extracellular vesicles (EVs) are small membrane-bound particles secreted by cells into the extracellular space (van Niel et al., 2018; Wang et al., 2024). They carry a wide range of biomolecules, including proteins, lipids, nucleic acids, and metabolites, that can be transferred between cells and influence various physiological processes. The content of EVs can vary depending on the cell type, the environmental conditions, and the state of the cell. As an organism ages, changes in the content, function, and fate of EVs can have significant implications for aging-related processes, including stress responses, inflammation, and tissue maintenance (Guix 2020; Manni et al., 2023). They play a key role in cell-to-cell communication and in maintaining proteostasis across different tissues (Beer and Wehman 2017; Wang and Barr 2018; Sarkar and Patranabis 2024). EVs can facilitate the exchange of protein quality control related molecules such as molecular chaperones, protein aggregates, signalling proteins between tissues, helping coordinate responses to cellular stress, damage, or misfolded proteins across the organism (Takeuchi et al., 2015). In the context of aging or diseases like neurodegeneration, EVs facilitate the transmission of both beneficial and harmful signals such as misfolded proteins (e.g., tau, amyloid-β) between cells, influencing the systemic response to stress, and potentially spreading pathological proteins across tissues (Wallis et al., 2020).

The content of the extracellular proteins and how do they change with age is of particular interest in the context of age-related diseases. Further how proteins involved in proteostasis and stress responses change with age help in understanding organism-wide proteostasis. Here we report a meta-analysis of EV proteins at organism level using *C. elegans* proteome data during conditions of aging, disease, stress and infection. We extended our analysis to protein abundance changes and aggregation with age in wildtype and also in lifespan mutant *C. elegans*. In addition, we also made comparison with EV proteome from human brain tissues affected with Alzheimer’s disease. Our analysis suggests that the EV associated proteins undergo significant changes during aging, disease as well as stress conditions. Aggregation of these proteins might affect EVs biogenesis, transport and their function offering an opportunity to identify potential biomarkers of age-related diseases.

## 2. Methods

### 2.1. Data collection and analysis

The EV proteome of *C. elegans* were downloaded from two published datasets (Russell et al., 2020; Nikonorova et al., 2022). The aging proteome of *C. elegans* was downloaded from published dataset (Walther et al., 2015). The aggregation or insoluble proteome datasets of *C. elegans* were downloaded from four published studies (David et al., 2010; Rodrigues et al., 2011; Walther et al., 2015; Anderton et a., 2024). The *C. elegans* heat shock datasets were downloaded from Larance et al., 2011 and the pathogen infection related dataset was downloaded from Treitz et al., 2015. The EV proteome data from human brain tissues was downloaded from Muraoka et al., 2020. The data was analysed in MS-Excel and using Perseus_2.0.6.0 program (Tyanova et al., 2016).

From the Nikonorova dataset, they covered a large number of EV proteins by isolating and separated according to heavy and light subunits. There are 7 replicates for light fractionation and 6 for the heavy one. We summed all the replicates and get rid of the proteins which have zero values (didn’t found in any replicate). After that, we got 2363 proteins in light subunit and 2277 in the heavy subunit, in which 1714 proteins to be common in both fractionations. In Russell dataset, we got 617 proteins which considered for further analysis. We merged the both datasets and got 3207 proteins after removing the duplicate proteins and used this dataset and our working EV protein set.

Gene Ontology (GO) term enrichment analysis of biological processes (BP) and cellular components (CC) was performed using Shiny GO 0.82 webserver (Ge et al., 2020) and Metascape (Zhou et al., 2019). Protein abundance changes in wildtype and lifespan mutant worms (*daf-2* and *daf-16* mutant worms) were calculated as fold changes at indicated time points (days) and comparisons were made relative to day1 or as mentioned in the results section.

### 2.2. Statistical analysis

Origin and GraphPad software were used for graphing and statistical analysis. Mann Whitney U test and statistical tests were done and p-values are mentioned in the relevant figure legends.

## 3. Results

### 3.1. Analysis of extracellular vesicles associated proteins

Extracellular vesicles (EVs) have been isolated and their content identified from various systems and tissues. They carry a spectrum of biomolecules such as proteins, lipids, RNA and metabolites and implicated in various cellular functions including as signalling molecules. Identification of EV proteome from a complete organism help in finding their role and possible molecular mechanisms of inter-tissue communication and organism wide proteostasis (Wang and Barr 2018; Guix 2020; Manni et al., 2023). To do this, we selected *C. elegans* model organism which is a good system to study aging, proteostasis and inter-tissue communication. There were two studies (Russel et al., 2020; Nikonorova et al., 2022) that reported isolation and identification of EV proteins from *C. elegans*. While Russel et al reported 513 proteins, Nikonorova et al reported 2841 proteins in the EVs (at least in one replicate). There were 348 common proteins (Figure 1A). We combined these two data sets to make a comprehensive list of 3207 EV proteins (Supplementary Table 1). We call these proteins as EVAPs (extracellular vesicles associated proteins). The EVs with their content leave the originating cells and travel in the extracellular fluid to reach their target cells. How do the EVAPs maintain their folded structure and function? To answer this, we checked whether the EVs also contain proteins related to proteostasis network (PN). We compared the known 821 PN components from *C. elegans* (Walther et al., 2015) with the EVAPs. We found that 250 proteins related to PN are associated with the EVs (Figure 1B). Among these 95 belong to protein synthesis, 89 belong to protein folding and 66 belong to protein degradation (Figure 1C). Based on these numbers, we assume that at least one fourth of the all PN components might present in the EVs. The GO term analysis for cellular components (CC) revealed that these EVAPs are part of vesicles, endomembrane systems and ribosomes (top 20 CC terms) and involve in localization, transport and metabolism related biological processes (BP) (Figure 1 D-E). Nikonorova et al also reported proteins from two types of EVs (heavy and light). Our GO term analysis revealed that there is some difference in the content of these two types of EVs (Suppl Figure 1). These finding confirm earlier observation that PN components can transmit via EVs to maintain organism-wide proteostasis (Takeuchi et al., 2015). In summary we made a comprehensive list of EVAPs from *C. elegans* model organism and found that they carry PN components in addition to many other classes of proteins.

**Fig. 1.**
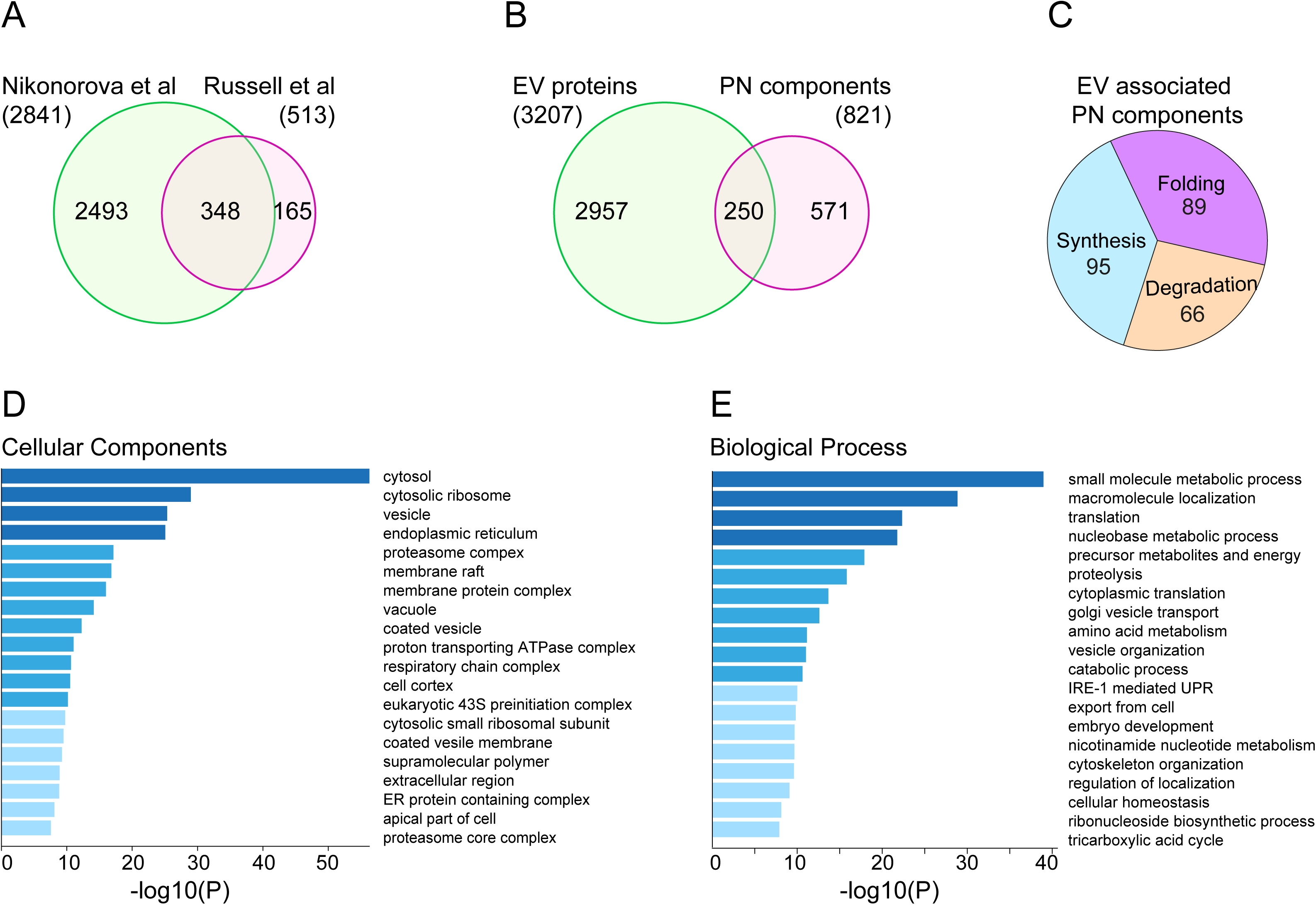
Identification of extracellular vesicles associated proteins (EVAP). **A.** Venn diagram showing the overlap between EVAPs found in light and heavy subunit of Nikonorova study with Russell study. There are 2841 and 513 proteins are found in Nikonorova both submit and Russell dataset respectively, in which 348 proteins are common in them. **B** Identification of the PN components in the EVAPs. There are 3207 unique EVAPs found from the two different EV studies and 821 PN component proteins found from the Walther study. And there 250 proteins are common between them. **C** The PN components associated with EVAPs. Out 250 EVAPs which were common with the PN components, 95 proteins are involved in synthesis, 89 in folding and 66in degradation, which are the three major arms of proteostasis. **D-E** Enriched gene ontology (GO) cluster of biological processes (BP) and cellular components (CC) of EVs found in both subunits respectively.

### 3.2. Identification of age-related changes in the EV proteins

Extracellular vesicles are implicated in aging and proposed to consider as hallmarks of aging (Yin et al., 2021; Manni et al., 2023; Bhunia et al., 2024). EV proteins were analysed from a few senescence cells or plasma from young and old animals. It has been reported that cellular senescence or aging alter EVs cargo (Alibhai et al., 2020). Further it has been observed that the EVs from young animals can improve health span of old animals (Sanz-Ros et al., 2022; Chen et al., 2024). However, many of these studies mainly focused on the miRNAs of the EVs and studies on proteins were limited.

To found how the EVAPs change with age in an organism level, we compared the EVAPs with an age-related proteome data by Walther et al 2015. In this study Walther et al., identified and quantified proteome wide changes during aging of wildtype and lifespan mutants (*daf-2* and *daf-16*) of *C. elegans*. We analysed abundance changes of the EVAPs in wildtype and lifespan mutants relative to day1 adult worms. We found significant change in the mean abundance levels with age in the wildtype as well as in the lifespan mutants (Figure 2A-C). These results suggest that the abundance of the EVAPs change in both directions. Then we specifically focused on proteins that increase or decrease more than 4-fold. We found that the EVAPs that increase more than 4-fold continue to increase with age. In wildtype and lifespan mutant worms, the abundance increase is more than the decrease (Figure 2 D-F). To gain more insight on the EVAPs that altered during aging, we selected proteins that either increase or decrease 2-fold at day 12 and performed GO term analysis (Suppl Figure 2 A-B). We only found minor changes in the biological process and cellular components among the strains. A more detailed analysis is required find changes that vary in the lifespan mutants. It is noteworthy that these protein abundance changes reflect at organismal level, not necessarily in the EVAPs. However, one can assume similar changes in the EVAPs as well because the proteomes were analysed by lysing whole worms.

**Fig. 2.**
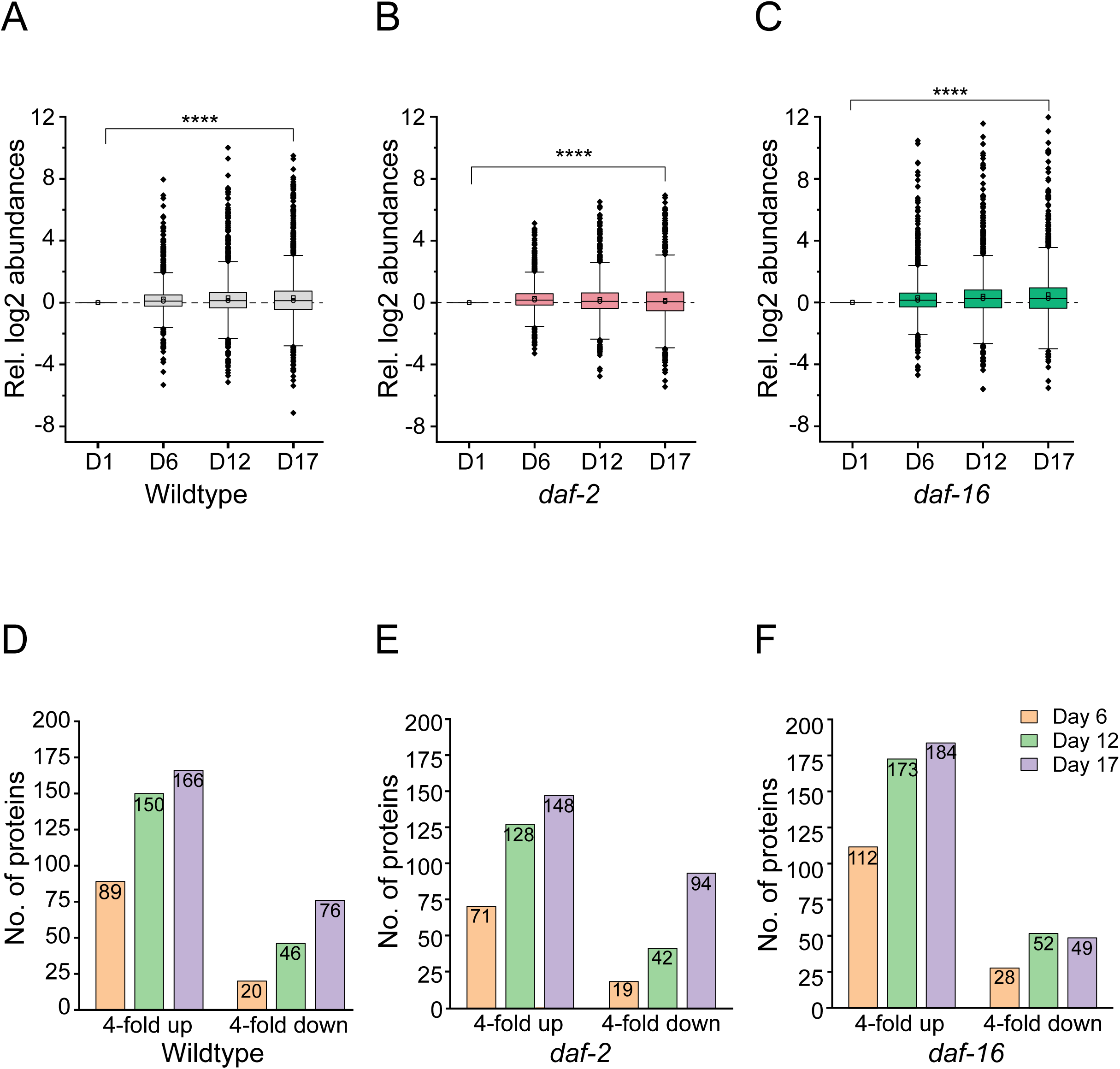
Abundance analysis of the EV proteins during aging. **A-C.** Boxplots showing the relative abundance of the EV associated proteins in Walther et al total proteome data in wildtype and lifespan mutant worms. The relative abundance level is calculated with respect to day1 values (plotted on the y-axis). Time points in days and strains are mentioned on the x-axis. (P-values: * *<*0.05, \*\**<*0.01, \*\*\**<*0.001, \*\*\*\**<*0.0001). **D-F** Bar chart representing the number of up and downregulated EV proteins (greater than 4-fold) that change with age in the Walther dataset. The different color codes for different days in WT, daf-2 and daf-16 mutants respectively. Number of proteins are plotted on the y-axis and time points in days are mentioned on the x-axis.

### 3.3. Aggregation of extracellular vesicles associated proteins with age

It has been reported that the EVs also carry misfolded, disease related proteins and implicated in age-related neurodegeneration (Hill 2019; Takeuchi and Nagai 2022). For example, Aβ peptides related to Alzheimer’s disease has been shown to interact with the EVs (Rajendran et al., 2006; Coughlan et al., 2024). Further it has been observed that the EVs can spread a disease as well as can cause diseases in healthy animals (Aulston et al., 2019; Shippey et al., 2022). Here we wanted to identify whether the EVAPs become insoluble or aggregated during aging. For this we analysed age-related proteome aggregation data from 3 different studies (David et al., 2010; Reis-Rudrigues et al., 2011, Walther et al., 2015). These studies reported aggregation of proteins during aging of *C. elegans* (day10 to day12 of adulthood) by different isolation methods. Walther et al study also reported aggregation data for the lifespan mutants (*daf-2* and *daf-16* mutant worms). Reis-Rudrigues et al identified 203, David et al identified 711 and Walther et al identified 1414 proteins that aggregated more than 1-fold in aged worms compared to day1 adult worms (Figure 3A). We merged these proteins and identified 1694 proteins that represent age-related insoluble or aggregated proteins (Supplementary Table 2). We found that 859 proteins that aggregate during aging overlap with the EVAPs which is corresponding to ∼65%. Recently, it has been reported that Amyloid β promotes protein aggregation similar to aging (Anderton et al., 2024). We wanted to check whether the Aβ induced protein aggregates are part of the EVAPs. We compared the Anderton et al data with the EVAPs and found that 452 of 595 proteins (∼76%) overlap with the EVAPs (Figure 3B-C). These observations suggest that the EVAPs prone to aggregation with age or disease conditions.

**Fig. 3.**
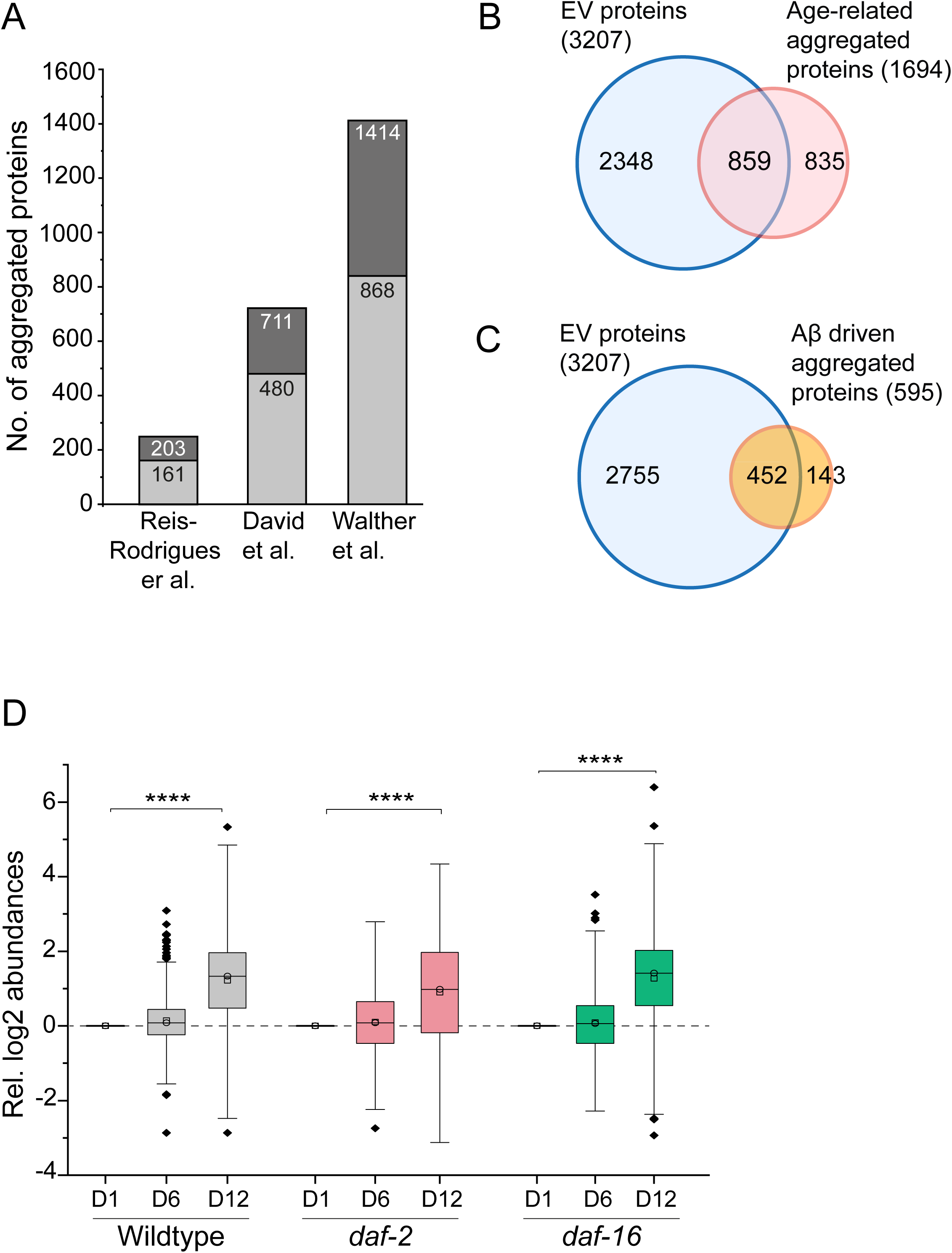
Insolubility of EVAPs during aging and disease condition **A** Bar chart representing the number of EV proteins found in aggregation datasets of Reis-Rodrigues, Della and Walther datasets respectively. Number of proteins are plotted on the y-axis and different datasets are mentioned on the x-axis. The deep grey color denotes the number of proteins found in the respective studies and the light grey color denotes the number of proteins overlapping with EV dataset. **B** Venn diagram illustrating the number of identified EV proteins in three different age-related aggregation datasets. There are 3207 EVAPs and 1694 age-related aggregated proteins, in which 859 proteins are common. **C** Venn diagrams showing the overlap between EVAPs with Aβ aggregation proteome dataset. In which 3207 are total EVAPs and 595 proteins are found in Aβ aggregation dataset and 452 proteins are between them. **D** Boxplots showing the relative abundance of the EV associated proteins in Walther et al aggregation data in wildtype and lifespan mutant worms. The relative abundance level is calculated with respect to day1 values (plotted on the y-axis). Time points in days and strains are mentioned on the x-axis. (P-values: \**<*0.05, \*\**<*0.01, \*\*\**<*0.001, \*\*\*\**<*0.0001).

Next, we looked at the protein aggregation during aging of the wildtype and lifespan mutant worms. Compared to the day1 worms, there is strong increase in the aggregation of the EVAPs with age in all the strains (Figure 3D). This is in line with the suggestion that the EVs can carry protein aggregates (Lim and Lee 2017). This suggest that either the proteins in the EVs aggregate during aging or the aggregated proteins pack into the EVs. We did not detect significant changes in the GO-terms among the worm strains (Suppl Figure 3). However, the long-lived *daf-2* mutant worms may properly maintain their proteins in the EVs. For example, the dauers, a specific larval stage shares many features with the *daf-2* long-lived mutant worms. Recently it has been reported that the EVs derived from the dauers extend lifespan of the wildtype worms (Ma et al., 2023).

In conclusion, we assume that the EVAPs aggregate during aging in *C. elegans*. Since the analysed proteome data were isolated by lysing whole animals, it is likely that the aggregated protein might represent both the intracellular and extracellular proteome including the EVAPs. Whether the aggregation of the EVAPs affect packing, formation or deliver of the EVs remain to be explored.

### 3.4. Association of proteins that alter during stress and infection with EV proteins

Extracellular vesicles are also implicated in protection from stresses such as heat stress and pathogen infections. EVs from heat stressed cells can promote survival of non-heat stressed cells by adaptive response, suggesting they carry protective molecules (Bewicke-Copley et al., 2017; Gebremedhn et al., 2020). During infections, EVs can also spread host as well as pathogen derived molecules to promote defence mechanisms and immunity (Rodrigues et al., 2018; Spencer et al., 2021). To identify whether the proteins that are altered during heat stress or pathogen infections are associated with the EVAPs, we analysed proteome data from *C. elegans* under heat stress (Larance et al., 2011) and pathogen infection conditions (Treitz et al., 2015).

First, we identified proteins that change their abundance during heat stress from both the data sets, combine them and sorted according to their fold change (more or less than 1.25-fold). We found 95 proteins upregulated and 484 proteins down regulated with heat stress. We found that 65 of the 95 upregulated proteins and 313 of the 484 downregulated proteins are associated with the EVs (Figure 4A). Next, we looked at the proteins that change their abundance with infection to the pathogen, Bacillus thuringiensis. We found 171 proteins upregulated and 117 proteins down regulated in response to the pathogenic bacteria. Out of the 171 upregulated proteins we found 110 proteins associated with EVs. In the downregulated proteins out of 117 we found 48 proteins were associated with EVs (Figure 4B). This analysis revealed that the EVs contain proteins that are required to respond to stress conditions such as heat stress or pathogen infections.

**Fig. 4.**
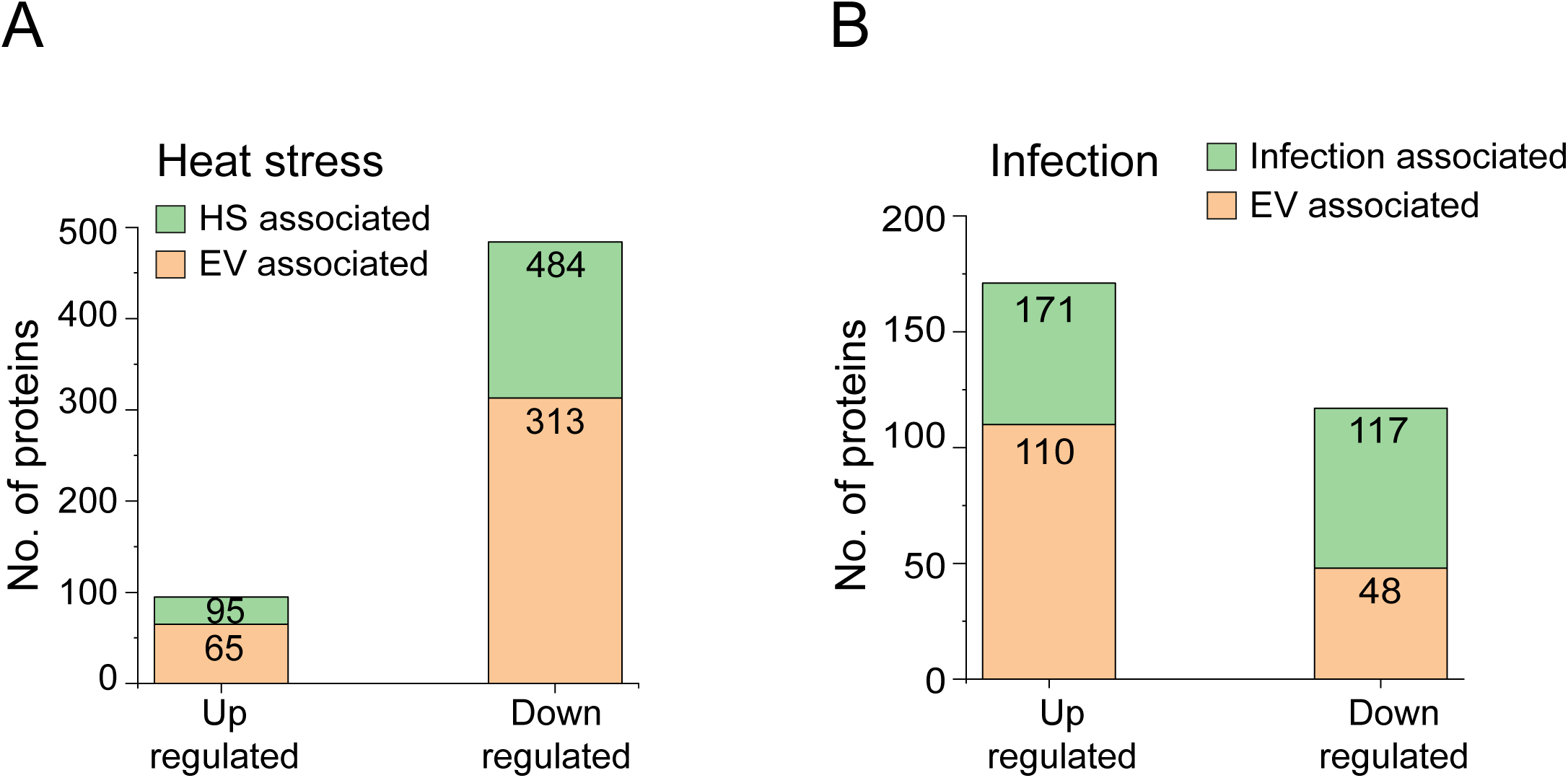
Overlap of EVAPs with the proteins that alter during heat stress and pathogen infection. **A** Bar chart representing the number of up and downregulated proteins during heat stress and the number of proteins among them which are EVAPs respectively. **B** Bar chart representing the number of up and downregulated proteins during pathogen infection and the number of proteins among them which are EVAPs respectively.

### 3.5. Overlap of C. elegans EVAPs with human EVs from brain tissues

EVs are implicated in age-related neurodegenerative diseases (Guix 2020; Yin et al., 2021; Xiao et al., 2021). EVs are involved in cellular communication as well as spread of pathology related to neurodegenerative diseases (Budnik et al., 2016; Sarkar and Patranabis 2024). The content of EVs determine their function. They function in protective manner by delivering neuroprotective factors or contribute to diseases by transferring toxic material (Tam et al., 2024). For example, EVs from brain tissues affect with Alzheimer’s disease (AD) carry amyloid plaques and tau tangles and can induce memory impairment in WT mouse (Bodart-Santos et al., 2023). This suggests EVs provide unique opportunity to identify for potential biomarkers for diagnosis and treatment of diseases (Zanirati et al., 2024; Pei et al., 2024). Towards this direction we wanted to check whether the EVAPs identified in *C. elegans* model organism have any human ortholog.

We analysed EV proteome data from human brain tissues affected by Alzheimer’s disease (Muraoka et al., 2020). A total of 949 proteins were quantified in this study, where 934 proteins are common between AD and control groups. Mean value fold change for control is 1.23 and 1.47 for AD which is statistically significant (Figure 5A). We also analysed log2 fold changes and found similar trend with average mean log2 values are −0.146 for control and −0.09 for AD (Suppl Figure 4A). We considered all the 949 EV derived proteins to identify human orthologs of *C. elegans* EVAPs and found 97 proteins (Figure 5B, (Supplementary Table 3). Next, we looked whether these human EVAPs also have any orthologs in the age-related or Aβ driven aggregated proteins. We found that 44 age-related and 22 Aβ driven aggregated proteins have human orthologs (Figure 5C-D). We further compared all the human orthologs and found significant overlap, where 15 proteins are common in all three conditions (Suppl Figure 4B). These observations suggest that a subset of the human brain derived proteins are prone to aggregation. To get a more insight, we checked fate of these 97 human orthologs during aging in wildtype and lifespan mutant worms. Intriguingly, we found that the total abundance of these proteins slightly increases with age in the wildtype worms. This abundance increase is more pronounced in the short-lived *daf-16* mutant worms. In contrast, the abundance of these proteins is less changed in the long-lived *daf-2* mutant worms (Figure 5E). Similarly, aggregation of these proteins is more prominent in the short-lived mutants compared to the long-lived mutant worms (Figure 5F). Next, we checked abundance of these proteins by comparing with all the identified proteins by label free quantification (LFQ) and found that the human orthologs have relatively more abundant in total as well as in the insoluble/aggregated proteome (Suppl Figure 4C-D). To know whether the *C. elegans* EVAPs or the human orthologs have intrinsic aggregation propensities to aggregate, we checked their aggregation propensities (fraction of insoluble protein of total protein). We found that many of these proteins have lower aggregation propensities (Suppl Figure 4E). This suggests that for majority of these proteins their increased abundance and supersaturation might result in their aggregation.

**Fig. 5.**
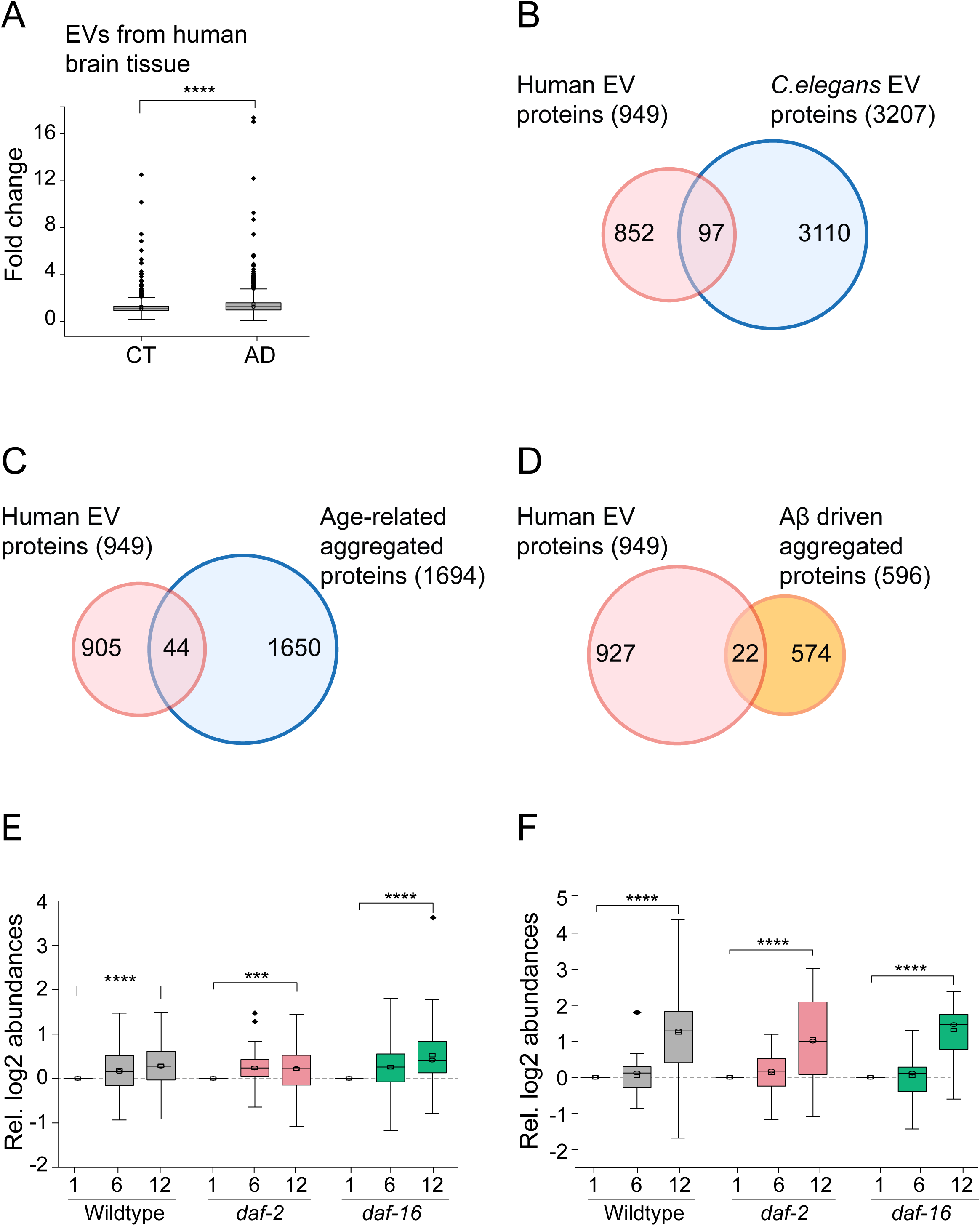
Overlap of *C. elegans* EVAPs with human EVAPs. **A** Boxplots showing the relative fold change of EVAPs in control and Alzheimer diseased group from human brain tissue samples. Mean value for control is 1.23 and 1.47 for AD. **B** Venn diagram showing overlap between human brain derived EVAPs and *C. elegans* EVAPs. 97 proteins have human orthologs. **C** Venn diagram showing overlap between human brain derived EVAPs and age-related aggregated proteins from *C. elegans* from three different studies. 44 proteins have human orthologs that present in the human EVs. **D** Venn diagram showing overlap between human brain derived EVAPs and Aβ driven aggregated proteins from *C. elegans*. 22 proteins have human orthologs that present in the human EVs. **E** Boxplots showing the relative total protein abundance of the human orthologs during aging in wildtype and lifespan mutant worms. The relative abundance level was calculated with respect to day1 values of each strain mentioned (plotted on the y-axis). Time points in days and strain names are mentioned on the x-axis. **F** Boxplots showing the relative protein aggregation abundance of the human orthologs during aging in wildtype and lifespan mutant worms. The relative abundance level is calculated with respect day1 values of each strain mentioned (plotted on the y-axis). Time points in days and strain names are mentioned on the x-axis. (P-values: * *<*0.05, \*\**<*0.01, \*\*\**<*0.001, \*\*\*\**<*0.0001).

## 4. Discussion

Inter-tissue communication has been observed across multiple tissues in a wide range of experimental systems and cell nonautonomous regulation of aging and stress response (O’Brien and van Oosten-Hawle 2016; Morimoto 2020). Organism-wide proteostasis involves complex cellular mechanisms that ensure protein quality control across different tissues (Sala et al., 2017). Inter-tissue communication, particularly via extracellular vesicles, plays a key role in maintaining organism-wide proteostasis by allowing for the exchange of signals and materials between cells and tissues. EVs are important players in this process, mediating both the spread of pathological proteins and potential protective responses that help maintain proteostasis at the organismal level (Takeuchi 2021; Wallis et al., 2020).

The fate of EV-associated proteins is of particular interest, as it remains unclear whether these proteins contribute to beneficial or harmful processes in recipient cells. Using the *C. elegans* model, we identified age-related changes in the composition of EV-associated proteins, including the aggregation of specific proteins and the accumulation of stress-induced factors, such as those triggered by heat stress and pathogen infection. These findings suggest that EVs may serve as a mechanism for the intercellular transfer of damaged or aggregated proteins, potentially contributing to the spread of cellular stress and the progressive decline in proteostasis observed during aging.

A key finding from this study is the identification of protein quality control (PQC) components as EVAPs, including chaperones, ribosome and degradation-related factors. The association of these proteins with EVs emphasizes their potential role in managing cellular stress and maintaining proteostasis through the intercellular transfer of quality-controlled proteins. However, with aging, we observed a shift in the nature of EVAPs, with a higher proportion of aggregated or misfolded proteins becoming associated with EVs. This suggests that as proteostasis deteriorates with age, EVs may become vehicles for the misdirected transfer of dysfunctional proteins.

Furthermore, the aggregation of EVAPs during aging has important implications for the biogenesis and delivery of EVs. The insolubility of aggregated proteins within EVs could interfere with their biogenesis, cargo sorting, or release, thus impairing the overall function of EVs. This disruption in EV function may further contribute to the decline in proteostasis and the progression of age-related diseases.

One of the most striking aspects of our analysis was the overlap between EVAPs and proteins that are known to be altered during heat stress and pathogen infection. We found that association of heat stress and pathogen infection-altered proteins with the EVs, which suggests that EVs play an important role in responding to environmental stressors and propagating stress responses across tissues. This association further underscores the dual role of EVs in both maintaining proteostasis and mediating cellular communication during stress conditions. The incorporation of misfolded or stress-related proteins into EVs may act as a protective mechanism to clear damaged proteins from the cells, but with aging, this process may become dysregulated, facilitating the spread of protein aggregation and exacerbating the aging phenotype.

We also found that orthologs of EVAPs from human brain tissues aggregate during aging and differentially regulated in lifespan mutant *C. elegans*. It has been observed that proteostasis is compromised in the short-lived mutant worms, while the long-lived mutants maintain properly balanced proteostasis. Presumably the EV content from the short-lived mutant worms might have more of misfolded proteins. Limitation of this study is that the data represents proteins from both intracellular and extracellular because of the use of whole worm lysate for proteomic identifications. Future studies focused on isolation and comparison of total as well as insoluble proteomes of EVs from young and aged samples as well as from lifespan mutant worms will be essential for further unravelling the complex role of EVs in aging and proteostasis.

## Acknowledgements

The authors acknowledge BioX centre, SBB at IIT Mandi for providing facilities.

## CRediT authorship contribution statement

Prasun Kumar: Data curation, Formal analysis, Investigation, Methodology, Validation, Visualization, Writing – review & editing. Prasad Kasturi: Conceptualization, Data curation, Formal analysis, Funding acquisition, Methodology, Supervision, Visualization, Writing – original draft, Writing – review & editing.

## Funding

This work was supported by funding from Department of Biotechnology, Government of India, as DBT-Ramalingaswami Re-entry grant (BT/RLF/Re-entry/31/2018 to P.K).

## Declaration of Competing Interest

The authors have no conflicts of interest to report.

## Appendix A. Supporting information

The supporting files containing datasets used in this work will be provided upon request.

---

**Supple Fig 1.**
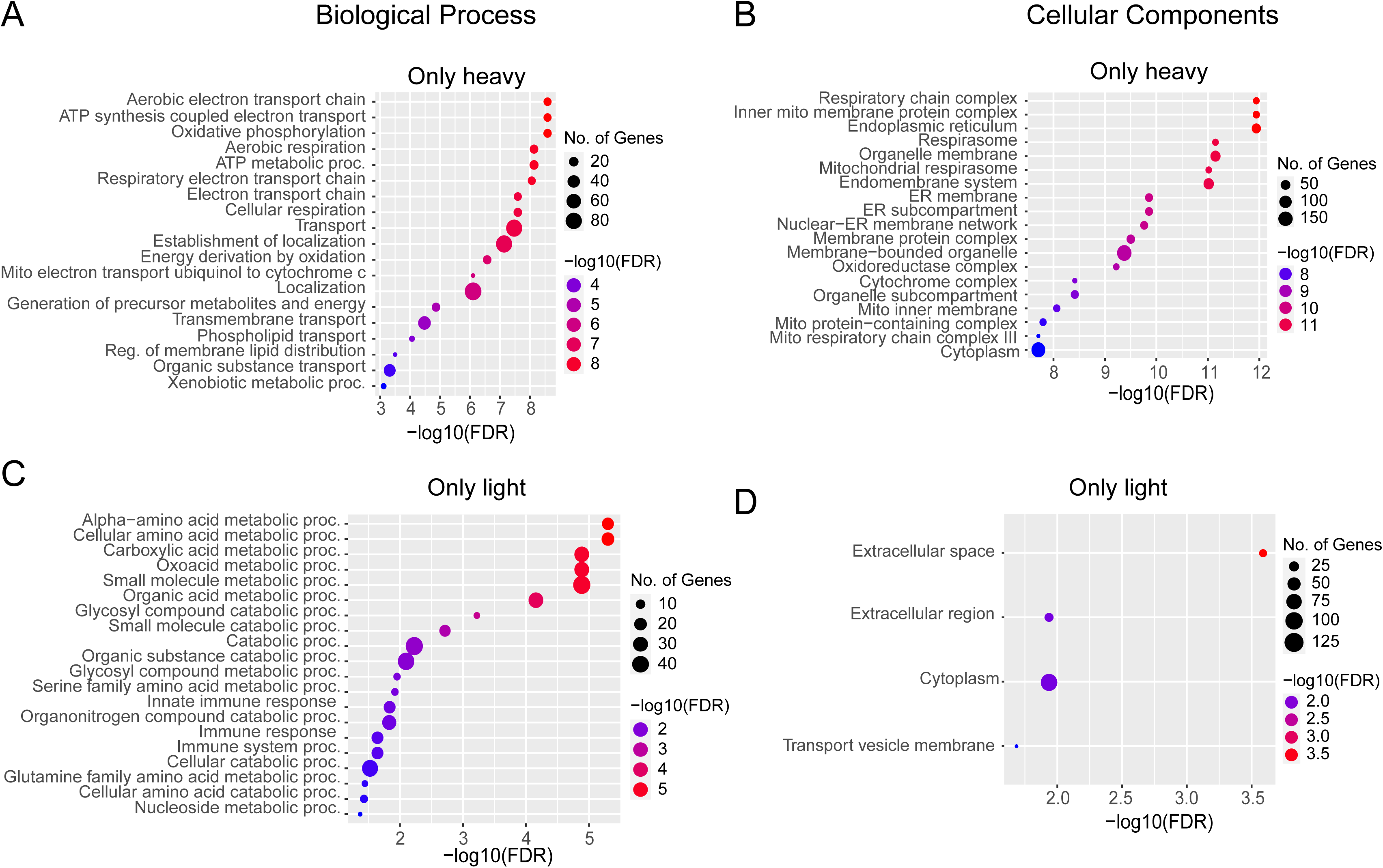
**A-B** Enriched gene ontology (GO) cluster of biological processes (BP) and cellular components (CC) of EVs found in only heavy subunit respectively. **C-D** Enriched gene ontology (GO) cluster of biological processes (BP) and cellular components (CC) of EVs found in only light subunit respectively.

**Supple Fig 2.**
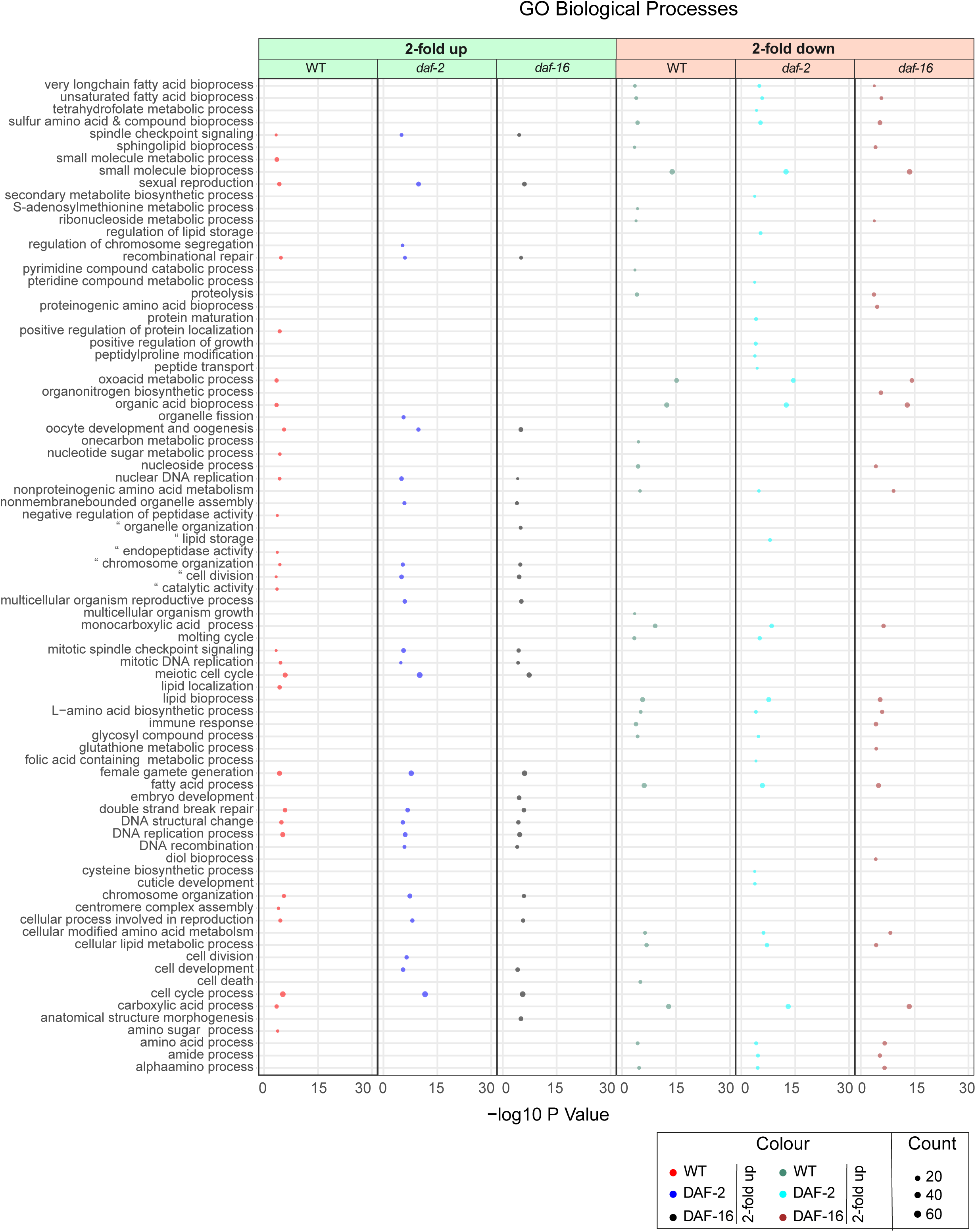
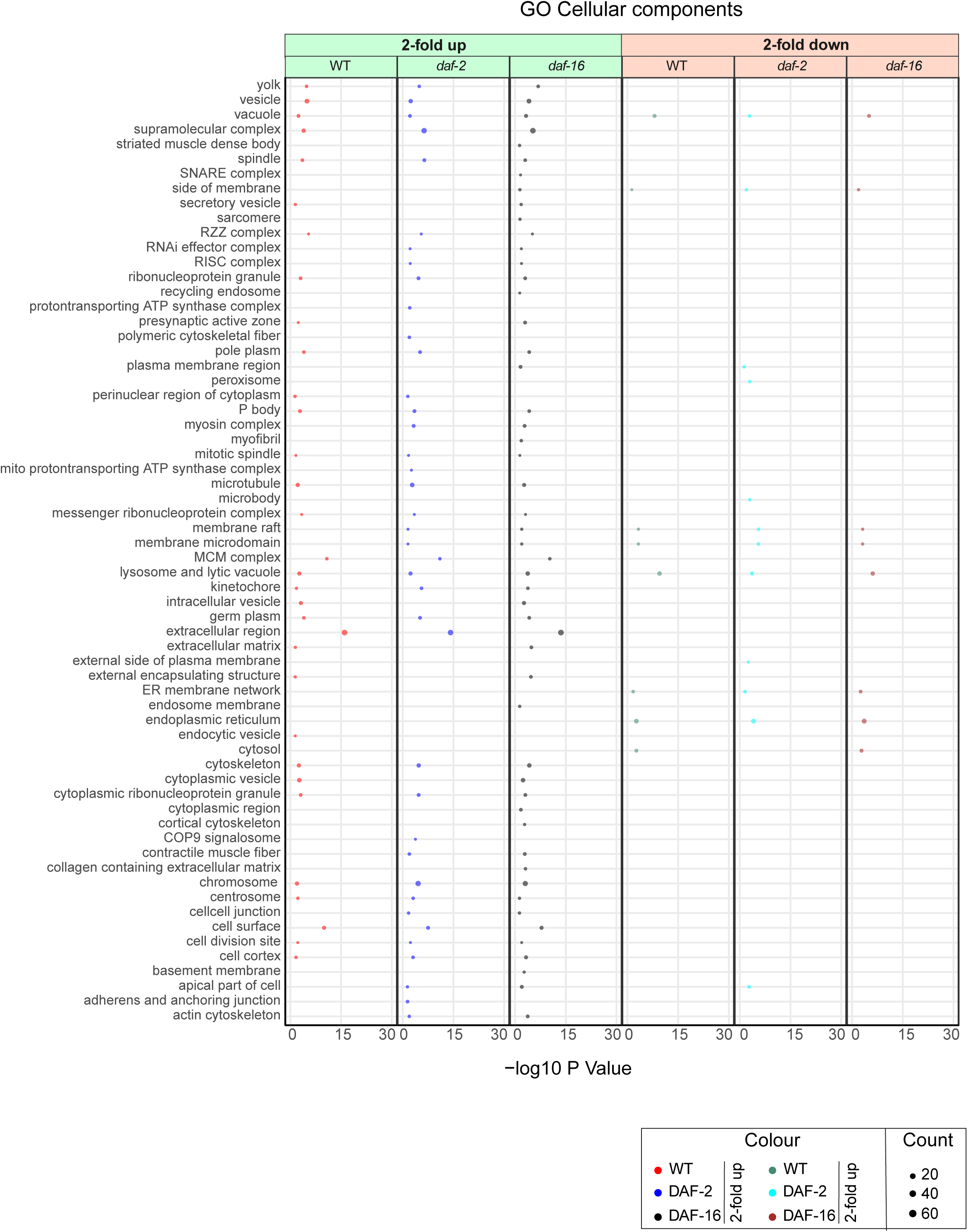
**A** Enriched gene ontology (GO) cluster of biological processes (BP) of the EVAPs upregulated and downregulated by 2 fold in day 12 in WT, *daf-2* and *daf-16* lifespan mutant worms. **B** Enriched gene ontology (GO) cluster of cellular components (CC) of the EVAPs upregulated and downregulated by 2-fold in day 12 in WT, *daf-2* and *daf-16* lifespan mutant worms.

**Suppl Fig 3.**
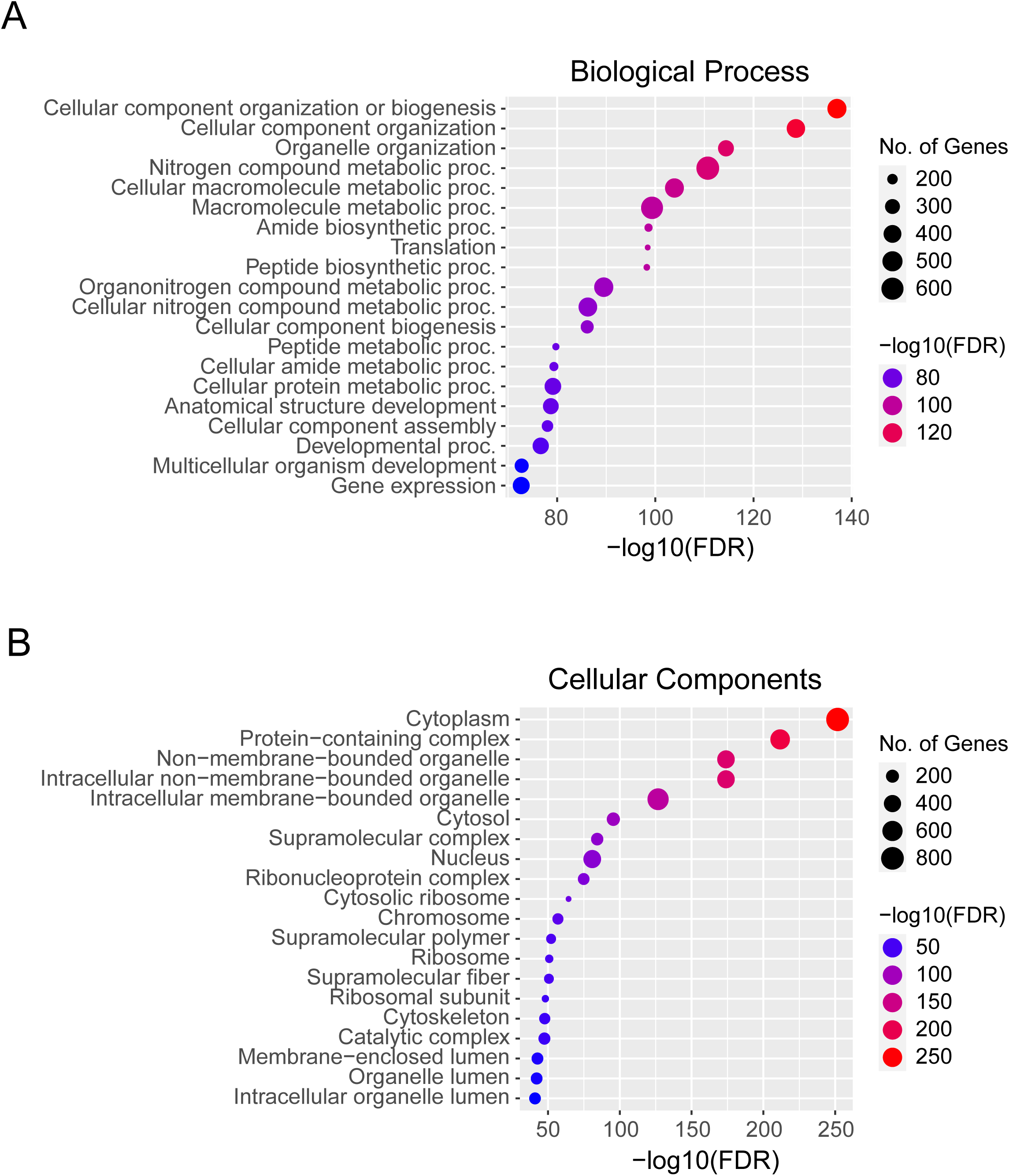
**A** Enriched gene ontology (GO) cluster of biological processes (BP) of the EVAPs that aggregate at day12 in wildtype worms. **B** Enriched gene ontology (GO) cluster of cellular components (CC) of the EVAPs that aggregate at day 12 in wildtype worms.

**Suppl Fig 4.**
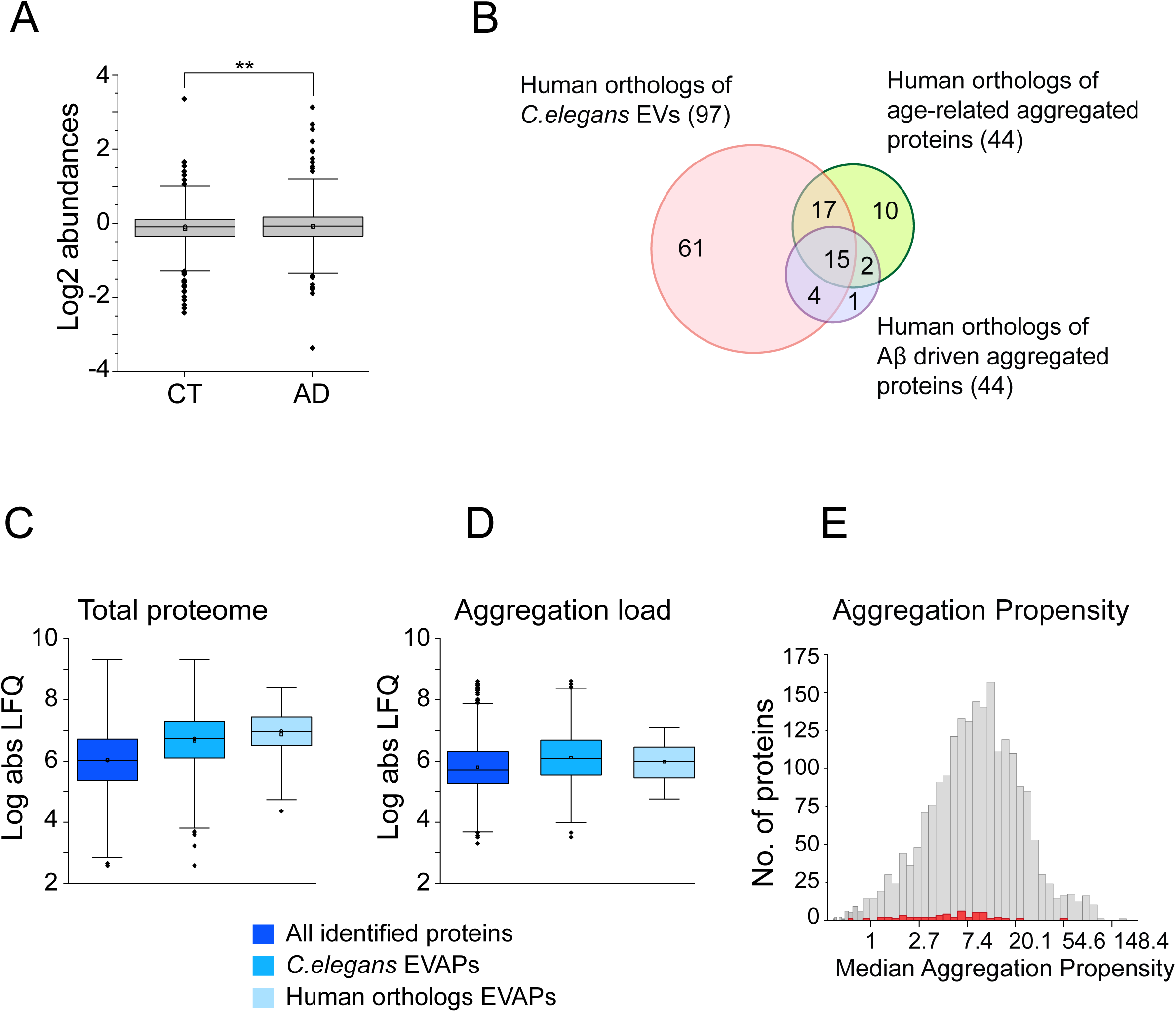
**A** Boxplots showing abundance (log2) of the proteins from control and AD human brain tissues. Average mean log2 values are −0.146 for control and −0.090 for AD. **B** Venn diagram showing overlap of human orthologs of EVAPs from *C. elegans* (97), age-related aggregated proteins (44) and Aβ driven aggregated proteins (22). 15 proteins are common in these three conditions. **C-D** Boxplots showing the relative abundance (log absolute LFQ) of proteins at day 12 in wildtype worms. **C** Total proteome abundance of all the identified proteins in Walther et al., study, *C. elegans* EV dataset and human orthologs. **D** Abundance of insoluble proteins of all the identified proteins in Walther et al., study, *C. elegans* EV dataset and human orthologs. **E** Histograms representing aggregation propensities. All identified proteins from Walther study shown in grey and the human orthologs of EVAPs are in red. from human studies. The x-axis represents median aggregation propensity percentage and y-axis the number of proteins.

## Notes

### Competing Interest Statement

The authors have declared no competing interest.

### Summary of Updates

A new section 3.5 is added; Figures are revised.

